# Distress feeding of depredatory birds in Sunflower and Sorghum protected by bioacoustics

**DOI:** 10.1101/200097

**Authors:** S.S. Mahesh, V. Vasudeva Rao, G. Surender, D.A. Kiran kumar, K. Swamy

## Abstract

One of the most ignored aspects of bioacoustic technology employed worldwide is lack of understanding between acclimatisation and distress feeding by depredatory birds. Acclimatisation results in gradual increase in resistance to bioacoustics in comparison to distress feeding, which makes sudden surge in instances of feeding by depredatory birds. Acclimatisation and distress feeding are independent functions of feeding behaviour. Distress feeding in itself is a function of physiological conditions of bird, extent of cropped area, distance traveled to obtain food, population dynamics, other natural habitats and cropping pattern in an area and is greatly influenced by them. There are no studies conducted to understand the distress feeding of birds in agricultural landscape. Experiments proved that bioacoustics could offer protection against distress feeding by birds although at reduced efficiency.

## Introduction

Bioacoustics like alarm, distress and predatory bird calls is chiefly used for protection of crops from bird depredation (Philip. C. Whitford, 2009). However, any bird management technique in agriculture, bioacoustics is bound to become less effective over a period of time, as the target species tend to ignore the threat indicated in the sounds. Birds tend to ignore alarm, distress and predator sounds if the same call or call sequence is played frequently (Fitzgerald, 2013). But birds tend to forcefully feed on the protected crops despite perceived dangers, when there are no alternate food source available for feeding and survival. Distress feeding has a characteristic of random occurrence and is different from acclimatisation which is a gradual process of adjusting to a perceived danger.

Distress feeding in depredatory birds can happen when there is a dearth of food in the surrounding areas and or when birds are breeding. The parent birds feed chicks (during *kharif*) at any cost, for it involves survival of the brood. Birds also tend to ignore the danger perceived at the protected fields when their density per unit area increases more than the supporting capability. Sometimes injured or maimed birds that cannot leave their roost in search of food, venture on distress feeding of nearby protected crops (*pers comm*. Surender and Swamy 2014).

The behavioural differences of parakeets such as movement between and within the habitats happens in relation to the availability of preferred food (Greene 1988). Distress feeding is a function of this. Damage to crop varies spatially and temporally owing to interactions between bird behaviour and population dynamics and crop type, location and phenology. The most common pattern is greater damage at the edge of the fields, decreasing with distance into the field interior. Damage to field interiors may also occur sporadically when flocks descend on fields. The distance that birds forage into the fields from the field edge may be influenced by the field layout, landscape surrounding the field, habitat affinity and distance to preferred habitat, food availability (within the field and more broadly), predation risk, escape behaviour, foraging behaviour and food gathering economics for birds (Institute for Land, Water and Society. 2013)

There are no studies done to check whether the distress feeding of protected fields by depredatory birds like Rose-ringed Parakeets and Baya Weavers occur or not. In order to check whether the birds ignore perceived dangers in a protected field, experiments were done in Sorghum and Sunflower fields to assess their behavioural pattern, extent of damage and record the reasons for such behaviour.

## Materials and Methods

Experiments were conducted at five locations viz., Sira (Karnataka) and Jukal, ICRISAT, ICAR-Indian Institute of Millet Research (IIMR) and Baswapur (Telangana). Sunflower was grown in Sira, ICRISAT and Baswapur, whereas, Sorghum was grown in Jukal and IIMR.

Four bioacoustic call sequences were developed during the study period (2012-2014). Call sequences were constructed using alarm, distress and predator sounds recorded from fields. Various techniques and parameters were employed to build the call sequence. Initially, common method vogue in most prominent international brands of bioacoustic equipments was tried (Hughes & Hughes, 2017). Calls were placed one after another in horizontal layout (call sequence-1 & 2) and later multilayering of sounds (vertical layout) along with horizontal layout was tried (call sequence-3 & 4).

Call sequence-1: The call sequence-I was built in Nov 2012 on the model of Punjab Agricultural University (1975) and International norms by placing one call after another and giving a long silence period of 20 min. Three calls were placed one after another viz., Rock Pigeon distress, House Crow alarm, and Rose-ringed Parakeet alarm. The total duration of construct-I was 43 min of which, calls occupied about 23 min. Of this, the longest call was of Rose-ringed Parakeet alarm (about 18 min). This construct was a moderate success in the fields of ICRISAT and ICAR-IIMR. The sequence gave protection against seven species of depredatory birds of agriculture.Call sequence-2: The experience gained from sequence-1 was put into use for building sequence-2. Long duration calls and silence was discarded as birds acclimatised quickly and fed during silence periods. The silence was broken into two slices (Figure-1) of 2 X 2.5 min. Rest 10 min in the call construct was occupied by sounds of House Crow distress, Common Myna alarm, Rose-ringed Parakeet alarm, predators like Shikra, Black Kite, Harriers and Falcons. Some of these calls were repeated to increase the length of sequence. The sequence gave protection against 12 species of depredatory birds of agriculture.

**Figure 1.**
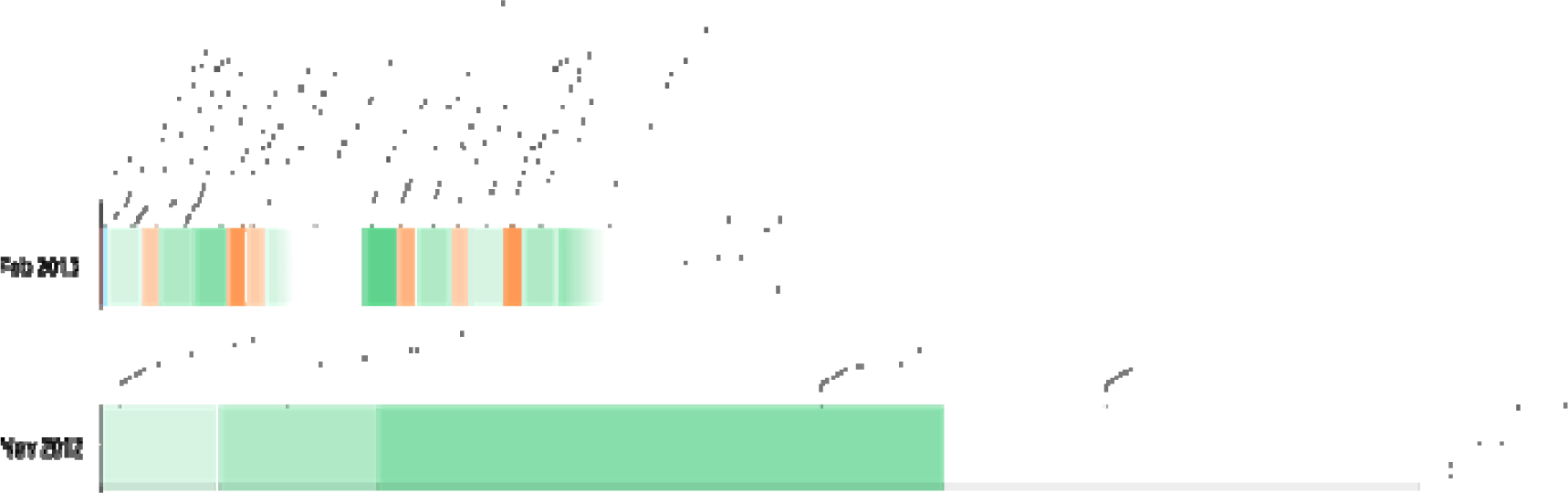
Structure of call sequence 1 & 2

Call sequence-3 contained bird sounds kept one over another (vertical layout), and one after another (horizontal layout) (Figure-2). It contains alarm complex of House Crow - Common Myna, Common Myna distress complex (repeats), Rock Pigeon chick distress, alarm calls of Indian Peafowl, Common Myna - Rosy Starling distress complex (repeats) and predator calls of Shikra and Black Kite. This call sequence also included artificial sounds like gun fire, and non avian bioacoustics like human shouts. Total duration of the sequence is 14 min.

**Figure 2.**
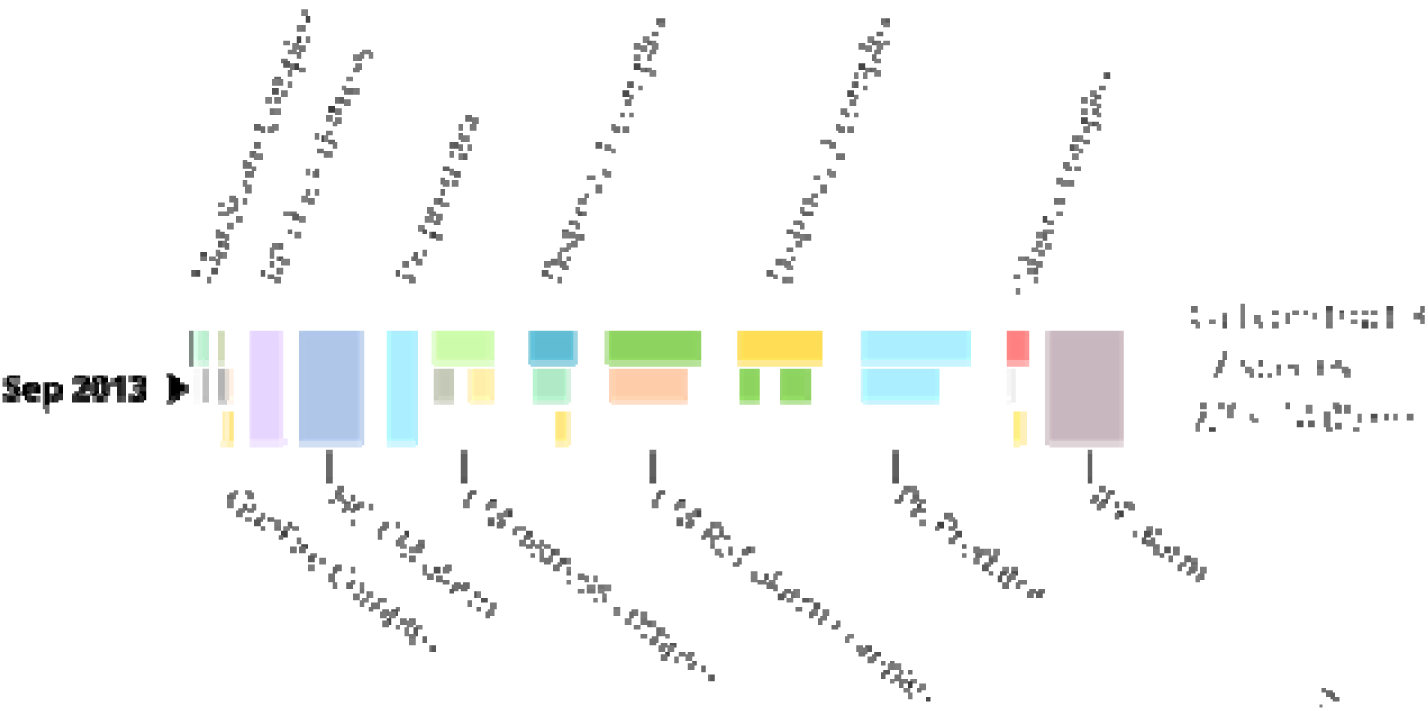
Structure of call sequence-3

An electronic broadcasting platform was assembled (Figure-3) with call sequences injected on to an electronic chip. Two equipments were fabricated for use in experimental sites. Each equipment had four audio output and were connected to full range 4Ω 35 w 10 cm Visaton speakers with frequency range of 80 Hz to 20,000 Hz. The call sequence-2 and 3 had frequency range of 200 Hz to 19,300 Hz, thus ensuring speaker outputted all frequencies that was desired to be broadcasted. Amplification was set to a level where each speaker gave an output of 105 dB sound level at source using alpha weighting method. A single device covered approximately four acres of area.

**Figure 3.**
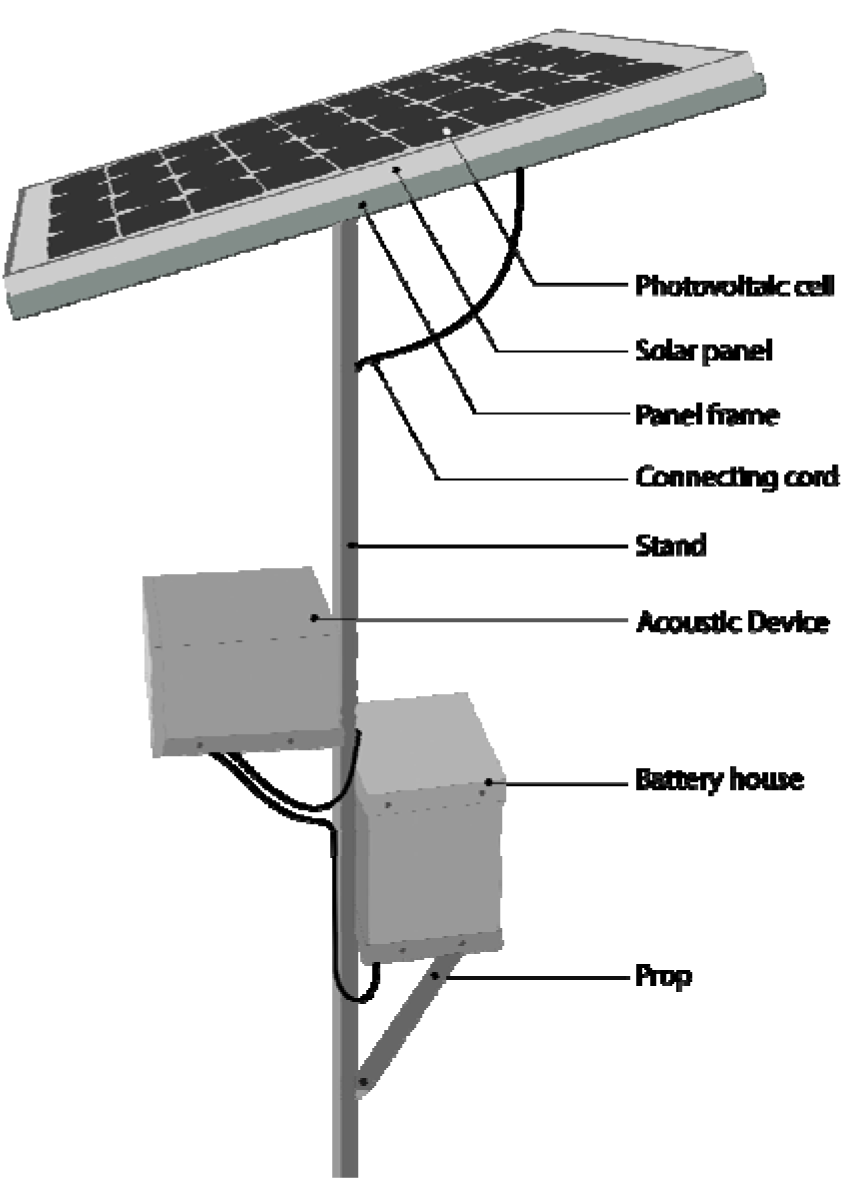
Bioacoustic Equipment Control unit

The speakers were installed at four corners of the field using extension cables. The height of the speakers was kept 30 cm above the crop canopy. The equipment derived its power from a 37 W 12 V solar panel and had a battery (12 V 26 Ah) as back up for 5-6 hours uninterrupted power supply. The equipment was turned on at sunrise and turned off at sunset. Bioacoustic equipment was removed after the crop was harvested and installed in another plot when the protection was needed. Two types of gaps between removal and installation of equipment was followed viz., 20 days and 90 days.

Time series observations were done from sunrise to sunset without a gap for the entire period of crop protection in experimental and control plots (Imadullah, 2014). Parameters like date, time, species, number visited, activity of the bird whether feeding, resting, perching or overflying etc), call played, and height were noted.

The efficacy of the bioacoustics was evaluated on 100 point Likert scale with differential weightings for depredatory behaviours. Four clearly differentiated behavioural pattern of Parakeets were recorded viz., Alert (initial response of the birds such as time taken to stop feeding, or look up, 10 marks), Lift (how hastily the birds took off and proportion of birds that took to wings, 10 marks), Hold (total time taken by birds to remain over the crop area, 10 marks) and Dispersal (total time taken by birds to disperse from the area of danger and proportion of birds that ultimately left the area of broadcasting, 70 marks) (Likert, 1932). In control plots, bird behavioural parameters were not included for observation as no bioacoustics was played there. In both the plots, initial damage if any, and damage on the day of harvest was calculated using standard techniques as prescribed by AINPAO (AINPAO, 2000).

The effectiveness of the bioacoustics was calculated by Inter quartile range (IQR) analysis where median score (50th percentile) was considered (Upton, 1996). Absolute dispersion for each season was calculated to know the reliability of equipment in dispersing parakeets. Comparison between IQR scores of different seasons were made to evaluate acclimatisation by birds to bioacoustics.

In all cases, experimental and control plots were of one acre and above, and away from each other at least a kilometre. Each experiment lasted for an average of 25-36 days from the beginning of formation of first achene of sunflower or grain of Sorghum to the day of harvesting. Details of location, crop, lat-long, size of the plot and general habitat features are given in (Table-1). All experiments were conducted in farmers’ fields. In the experimental plots, only bioacoustics was played as a means of bird management, whereas in control plots, occasional human shouting was carried out for dispersing depredatory birds.

**Table 1.** Location and crop details of experiments conducted

| Location | Duration | Crop | Lat-Long | Size of the plot (ha) | General habitat |
| --- | --- | --- | --- | --- | --- |
| Sira | Aug 2013 | Sunflower | N13.820026, E76.913626 | 0.4 | Predominantly agriculture intercepted by fallow lands and scrub jungle, Wetland areas. |
|  | Oct 2013 | Sunflower | N13.817467, E76.908281 | 0.4 |  |
|  | Jan 2014 | Sunflower | N13.810068, E76.882753 | 0.8 |  |
|  | Mar 2014 | Sunflower | N13.820074, E76.871136 | 0.4 |  |
|  | Aug 2014 | Sunflower | N13.820026, E76.913626 | 0.6 |  |
| Baswapur | Dec 2013 | Sunflower | N18.162735, E78.417175 | 0.5 | Predominantly agriculture |
|  | Mar 2014 | Sunflower | N18.143164, E78.404596 | 0.8 |  |
| IIMR, Hyderabad | Feb 2013 | Sorghum | N18.166798, E78.425415 | 0.4 | Urban infrastructure with fragmented agriculture holdings |
| ICRISAT | Jan 2013 | Sunflower | N17.515548 E78.267701 | 0.4 | Agriculture |
| Jukal, Hyderabad | Sep 2013 | Sorghum | N17.215950, E78.298900 | 0.8 | Predominantly agriculture, Scrub jungle |

Extent of cropped area was assessed by visiting the surrounding areas on regular intervals to know the availability of food to Parakeets and Baya Weavers. Land use land cover maps were used for assessing the extent of cropped area in the study locations. It is a known fact that Parakeets fly 2.5 to 17.25 km/day oneway for feeding depending on the availability of food (AINPAO, 2012). In the present study, we estimated flying distance of 5 km by Parakeets to access food, resulting in survey of 78 km^2^ of area. Roost studies of Parakeets were done in all locations of study, to understand the range of operation of Parakeets. The experimental and control plots in both the sites were less than four km from the roost. For breeding Baya Weavers, number of nesting birds was counted and the birds were followed on foot to know their extent of operation. The experimental site was less than 100 m from the breeding site. Similar method was adapted for breeding Rose-ringed Parakeets. Nesting colony of parakeets was 200 m from the experimental plots.

Distress feeding was studied using three approaches viz., switching off the bioacoustic equipment in between, assessing the cropping extent before the start of experiment and during surge in feeding instances, and breeding observations on Parakeets and Baya Weavers. Switching off the bioacoustic equipment in between was done only in one experiment viz., Aug 2014. In all other experiments, cropping extent and breeding observations were done to assess the distress feeding

## Results and discussion

Out of 10 experiments conducted, distress feeding happened in five instances viz., Oct 2013, Jan 2014, Aug 2014 (all in Sira) and Mar 2014 (in Baswapur) in Sunflower crop, and one instance of distress feeding was noted at IIMR during Feb 2013 in Sorghum crop. Call sequence-3 was used in all Sunflower experiments and call sequence-2 was used in Sorghum (Table-2). Distress feeding by Parakeets and Baya Weavers was noted during experiments done in Oct 2013. In all other experiments, only Parakeets performed distress feeding.

**Table 2.** Details of experiments

| Place of experiment | Season | Crop | Distress Feeding | Distress feeding by bird species | Call sequence |
| --- | --- | --- | --- | --- | --- |
| Sira | Aug 2013 | Sunflower | No | - | 2 |
|  | Oct 2013 | Sunflower | **Yes** | Rose-ringed Parakeet<br>Baya Weaver | 3 |
|  | Jan 2014 | Sunflower | **Yes** | Rose-ringed Parakeet | 3 |
|  | Mar 2014 | Sunflower | No | - | 3 |
|  | Aug 2014 | Sunflower | **Yes** | Rose-ringed Parakeet | 3 |
| Baswapur | Dec 2013 | Sunflower | No | - | 3 |
|  | Mar 2014 | Sunflower | **Yes** | Rose-ringed Parakeet | 3 |
| IIMR | Feb 2013 | Sorghum | **Yes** | Rose-ringed Parakeet | 2 |
| ICRISAT | Jan 2013 | Sunflower | No | - | 1 |
| Jukal | Sep 2013 | Sorghum | No | - | 3 |

## Sira Experiments

### Experiments of 12 Sep 2013 to 17 Oct 2013, Sira

Distress feeding by Parakeets happened at three of the five experimental locations conducted at Sira. During the experiments of Sep-Oct 2013, the plot was damaged by Parakeets to an extent of 0.05% (negligible damage) before installing the equipment (12 Sep 2013). Distress feeding by Parakeets and Baya Weavers started on 16^th^ day of installing the bioacoustic equipment (Figure-4). Distress feeding continued for 15 days till the crop was harvested on 36^th^ day (17 Oct 2013). The efficiency of bioacoustics in first 15 days was 85%. From 16^th^ to 26^th^ day, the efficiency of bioacoustics declined suddenly to 34%. In the last 10 days of experiment where distress feeding of Parakeets and Baya Weavers combined with acclimatisation to bioacoustics, the efficiency gradually declined to 22% till the crop was harvested.

**Figure 4.**
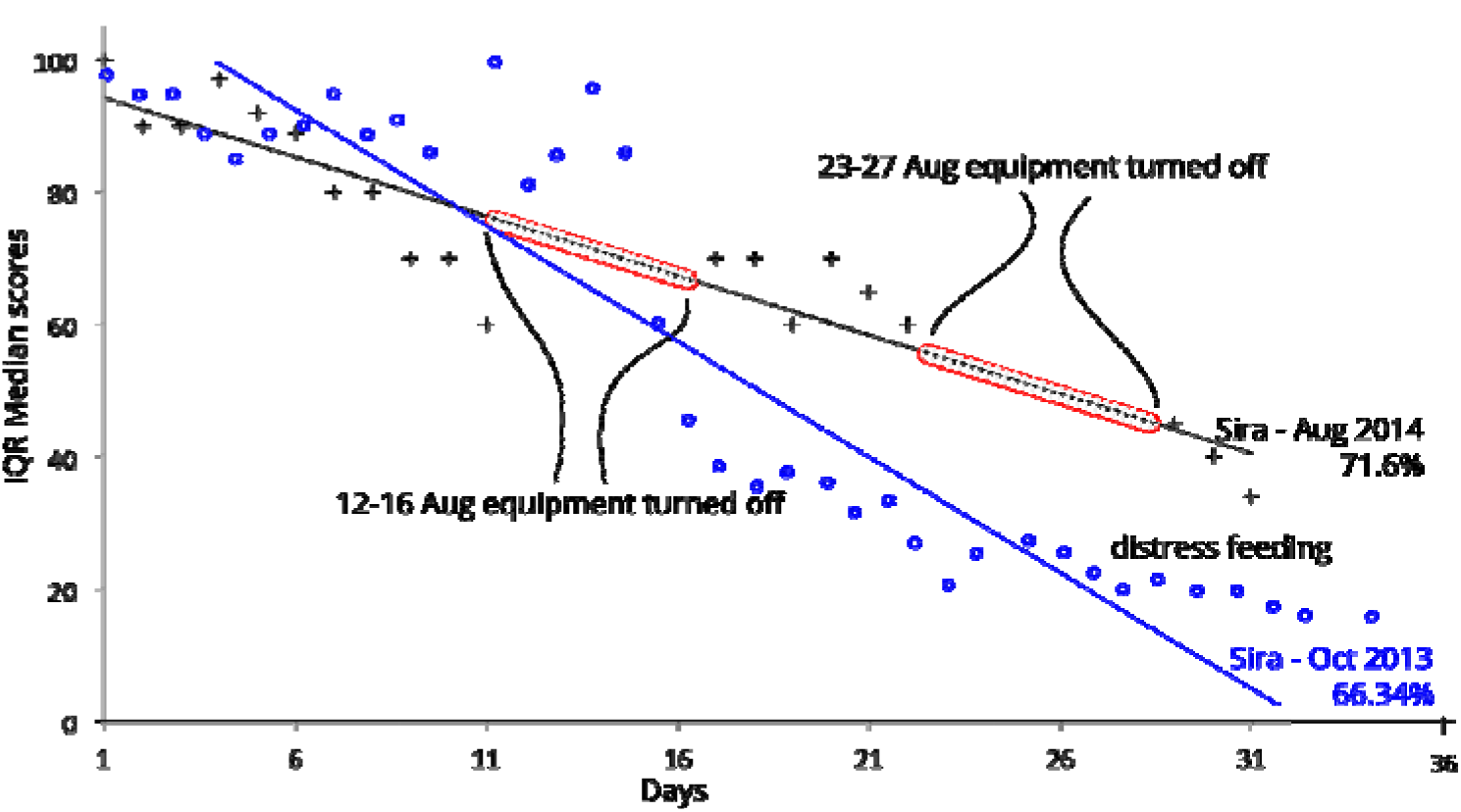
Distress feeding pattern by Parakeets and Baya Weavers

From the land use and land cover maps for this experimental locality, we found that only 60% (46.8 km^2^) of the area had plants and trees that were providing food to Parakeets and Baya Weavers. Of this cropping extent, an average 63% of plants/trees bore food in the form of flowers, young fruits, or seeds edible by depredatory birds. It is evident from (Table-3) that in the beginning of experiment availability of Sunflower, Ragi, Maize and Fodder Sorghum for Parakeets was plenty. However, on the date of distress feeding, availability of all of them had decreased considerably. For example most of the Sunflower crops were harvested by the end of experiment, only 28 ha of area was available. Parakeets preferred Sunflower over all other food at all points in time during the experiments. Though many of the highly preferred crops like Pomegranate were available, these crops were well guarded and farmers didn’t let Parakeets feed. Similarly for Baya Weavers, who were breeding during that time, the availability of food plants like Sunflower, Fodder Sorghum, Ragi, Grasses were declining. However, they completed breeding and abandoned nests by the end of Sep 2013. Thus their pressure on Sunflower in the end decreased.

**Table 3.** Extent of cropped area and availability of food to Parakeets and Baya Weavers, Sep-Oct 2013

| Crops with feeding stage for Parakeets and Bayas in 78 km<sup>2</sup> | Extent of cropped area that has potential to provide food (ha) | Extent of area providing food as on 12 Sept 2013: beginning of experiment (ha) | Extent of area providing food as on 06 Oct 2013: at the start of distress feeding (ha) | Extent of area providing food as on 17 Oct 13: at the harvest time (ha) |
| --- | --- | --- | --- | --- |
| Sunflower | 234 | 187 (80%) | 77 (33%) | 28 (12%) |
| Ragi | 1872 | 1479 (79%) | 280 (15%) | 131 (7%) |
| Maize | 374 | 250 (67%) | 82 (22%) | 7.5 (2%) |
| Fodder Sorghum | 936 | 749 (80%) | 47 (5%) | Nil |
| Grasses (in patches) | 234 | 210 (90%) | 187 (80%) | 145 (62%) |
| Pomegranate | 140 | 70 (50%) | 92 (65%) | 112 (80%) |
| Wild flowering and fruiting trees, Lantana etc | 486 | 194 (40%) | 233 (48%) | 267 (55%) |
| Other minor crops (Rice, Bajra, Pigeonpea, Mango) | 421 | 63 (15%) | 101 (24%) | 126 (30%) |

Visiting instances of Parakeets from the beginning of the experiment to 15^th^ day was 45/day. The instances of visits increased to 72/day from 16^th^ to 26^th^ day. After 26^th^ day till harvest, the visiting instances of Parakeets remained at 70/day. This showed that the surge in instance of Parakeets visiting experimental plots was correlated to shortage of food in the feeding range. The Parakeets sustained the visiting instances in the same tempo till harvesting. Similarly, visiting instances of Baya Weavers from the beginning of the experiment to 15^th^ day was 14/day. The instances of visit increased to 16/day from 16^th^ to 26th day. After 26^th^ day till harvest, the visiting instances of Baya Weavers remained at 6/day. There was no surge in instances of Baya Weavers visiting the field. This was mainly due to very limited and localised feeding behaviour of breeding birds, apart from absence of choice of food based on palatability by Bayas. Bioacoustics provided 85% protection for first 15 days in the absence of distress feeding, and continued to give protection, albeit in lesser efficiency, 73%, till 25 days. Overall efficiency further reduced to 66.3% for the entire crop protection period covering distress feeding and acclimatisation. Crop damage was negligible (1.06%) compared to 91% in control plot (Table-5).

### Experiment 17 Dec 2013 to 12 Jan 2014, Sira

During this experiment, the plot was damaged by Parakeets to an extent of 0.01% (negligible damage) before installing the equipment (17 Dec 2013). Distress feeding by Parakeets started on 21^st^ day of installing the bioacoustic equipment (Figure-5). Distress feeding happened for five days till the crop was harvested (12 Jan 2014). The efficiency of bioacoustics in first 21 days was 98%. From 22^nd^ to the end of experiment, the efficiency of bioacoustics declined suddenly to 14% till the crop was harvested.

**Figure 5.**
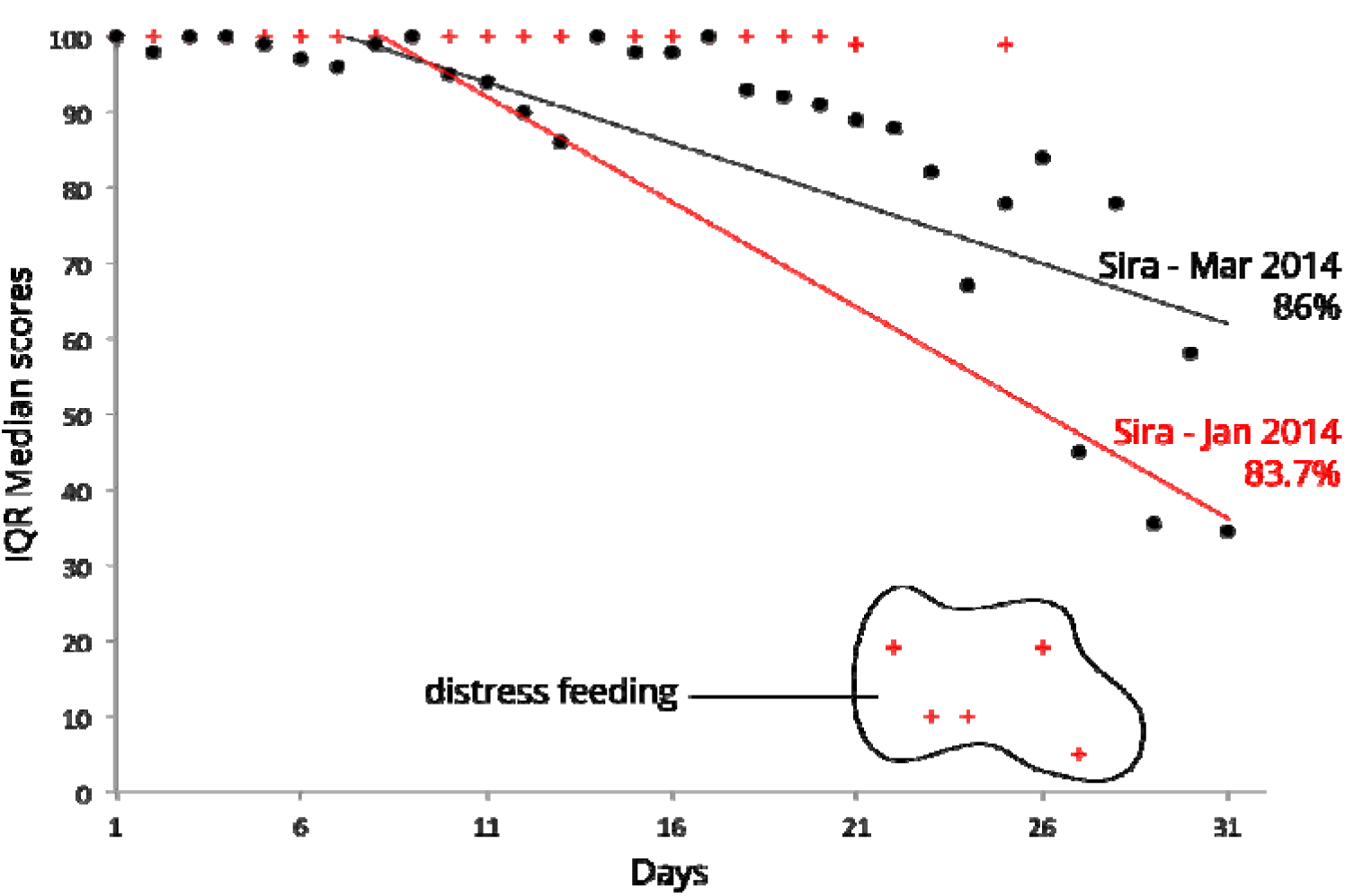
Efficacy of bioacoustics in Sunflower at Sira during Jan and Mar 2014

From the land use and land cover maps for this experimental locality, we found that only 48% (35.1 km^2^) of the area had crops and trees that were providing food to Parakeets. Of this cropping extent, an average 62% of crops/trees bore food. It is evident from (Table-4) that in the beginning of experiment, availability of Sunflower, Ragi, Maize, Pomegranate, wild flowers and minor crops for Parakeets was plenty. However, on the date of distress feeding, availability of all of them had decreased considerably.

**Table 4.** Extent of cropped area and availability of food to Parakeets, Dec 2013 to Jan 2014

| Crops with feeding stage for Parakeets in 78 km<sup>2</sup> | Extent of cropped area that has potential to provide food (ha) | Extent of area providing food as on 17 Dec 2013: beginning of experiment (ha) | Extent of area providing food as on 07 Jan 2014: at the start of distress feeding (ha) | Extent of area providing food as on 12 Jan 14: at the harvest time (ha) |
| --- | --- | --- | --- | --- |
| Sunflower | 246 | 185 (75%) | 89 (34%) | 81 (33%) |
| Ragi | 1229 | 959 (78%) | 221 (18%) | 221 (18%) |
| Maize | 421 | 337 (80%) | 101 (24%) | 97 (23%) |
| Pomegranate | 105 | 65 (62%) | 57 (54%) | 57 (54%) |
| Wild flowering and fruiting trees | 878 | 474 (54%) | 457 (52%) | 457 (52%) |
| Other minor crops (Rice, Pigeon pea, Mango) | 632 | 145 (23%) | 114 (18%) | 114 (18%) |

Visiting instances of Parakeets from the beginning of the experiment to 21^st^ day was 83/day. The instances of visits increased to 204/day from 22^nd^ day till harvest. This showed that the surge in instance of Parakeets visiting experimental plots was correlated to shortage of food in the feeding range. In this experiment, bioacoustics provided 98% protection for first 21 days in the absence of distress feeding, and continued to give protection, albeit in lesser efficiency, 14%, till crop was harvested. Overall efficiency further reduced to 83.7% for the entire crop protection period covering distress feeding. Crop damage was negligible (0.04%) compared to 43.8% in control plot (Table-5).

**Table 5.**
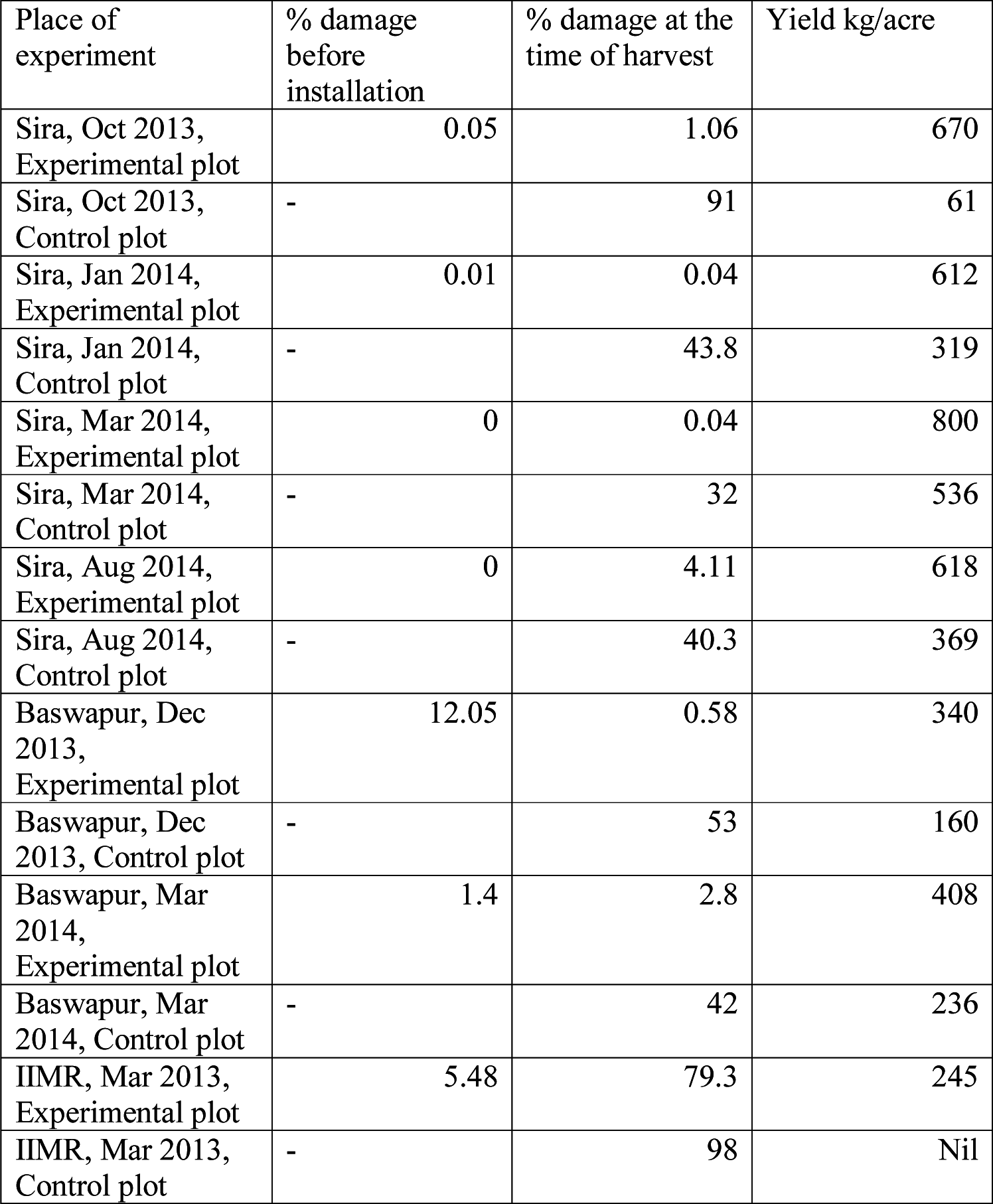
Details of experiment conducted along with damage percentage

### Experiment 01 Aug 2014 to 04 Sep 2014, Sira

There was no damage by Parakeets in experimental plot before installing the equipment (01 Aug 2014). Distress feeding by Parakeets started on 9^th^ day of installing the bioacoustic equipment (Figure-4). Distress feeding happened for three days till a decision to switch off bioacoustic equipment was taken. Bioacoustic equipment was switched off for four days twice (12-16 Aug 2014 and 23-27 Aug 2014) (Figure-4). During this period, a sudden surge in the number of instances of Parakeets visiting the plots was recorded (Figure-6). Up to ninth day of experimental period, the instances of Parakeets visiting Sunflower was ranging from 20-40 per day. On tenth and eleventh days, there was an indication of increase in instances (200-300 per day). It was then the equipment was switched off on 12th to 16th days. During this period, the instances of Parakeet visits increased to 1250 per day. On the17th day, equipment was turned on till 22nd day. During this period, the instances declined to an average of 230-300 instances per day coinciding with the instances of tenth and eleventh day. When equipment was again switched off on 23rd to 27th day of experiment, the instances surged again to 1500 per day coinciding with the instances of 12th to 16th day. From 27th day to 34th day, instances of Parakeets reduced to 30-200 per day. This showed that the Parakeets distress fed on Sunflower and did not acclimatise to bioacoustics. Crop was harvested on 35th day.

**Figure 6.**
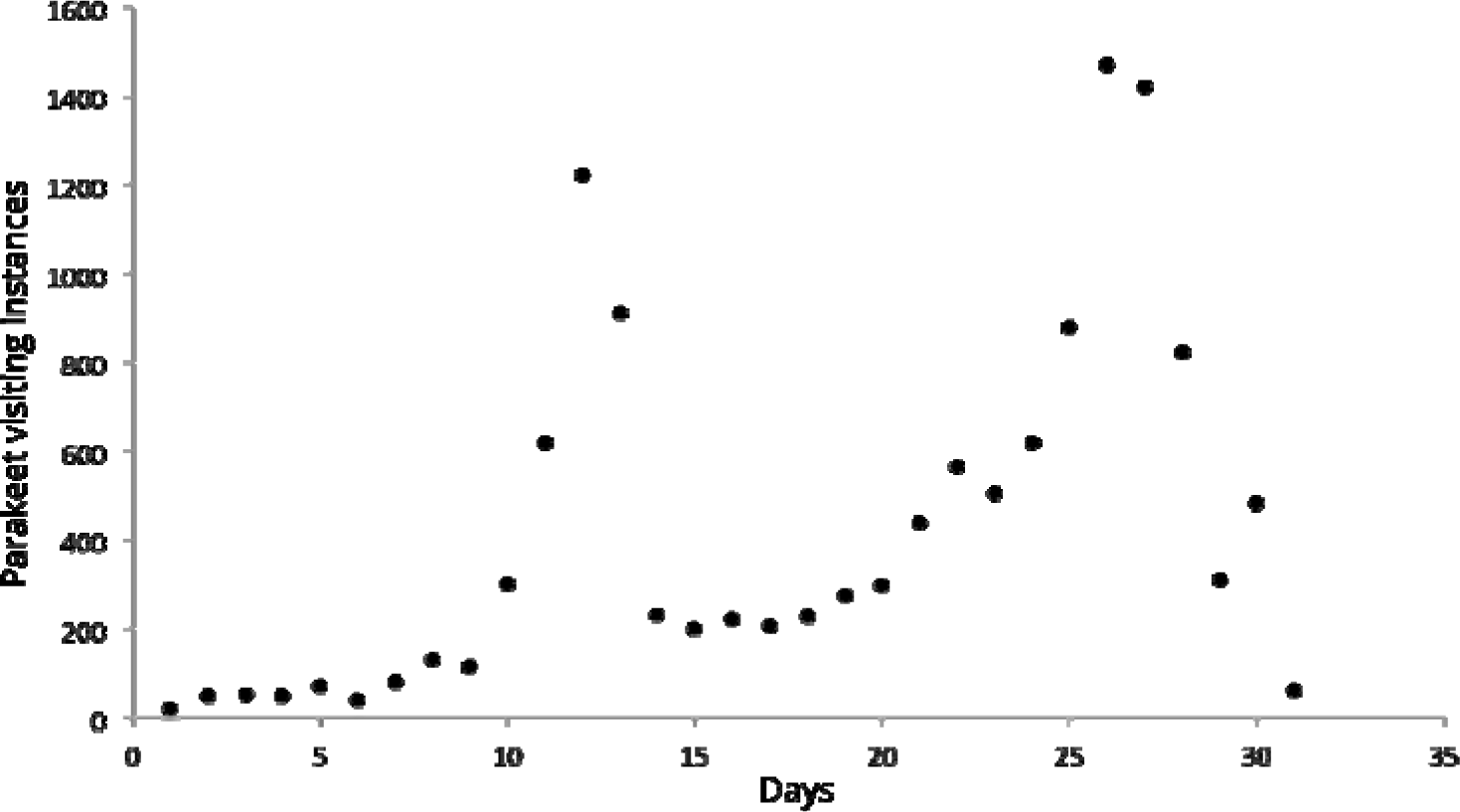
Surge in Parakeet visiting instances at experimental plot when bioacoustic equipment was switched off on 12-16 Aug & 23-27 Aug 2014

**Figure 7.**
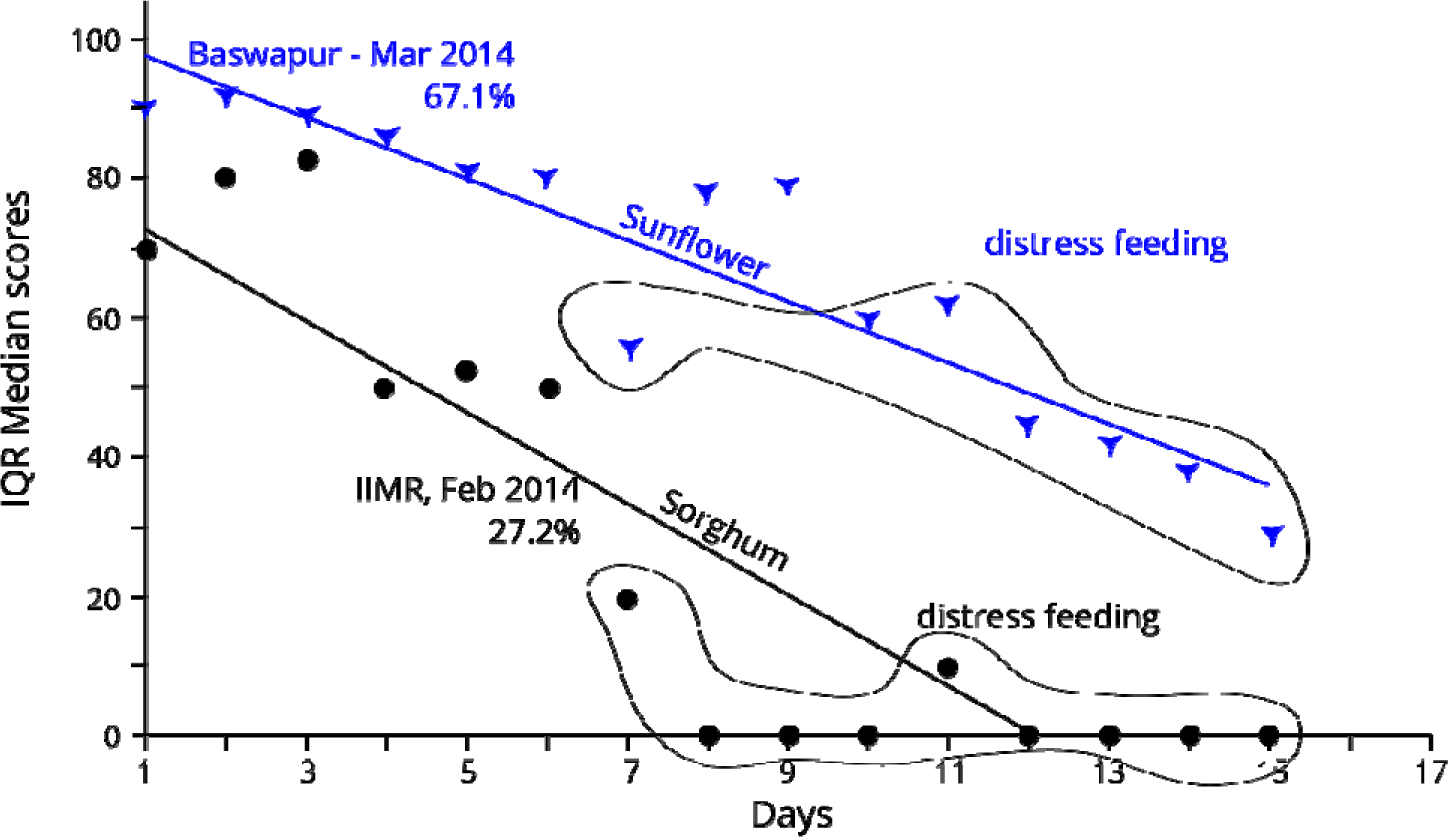
Efficacy of bioacoustics in Sunflower and Sorghum, Feb-Mar 2014

From the land use and cover maps for this experimental locality, we found that the parameters were similar to (Table-3). In the beginning of experiment, availability of food and minor crops for Parakeets was abundant. However, on the date of distress feeding, availability of food had decreased considerably.

The efficiency of bioacoustics in first eight days was 90.63%. From 9^th^ to 11^th^ day, the efficiency of bioacoustics dropped suddenly to 66%. When bioacoustics was reinstalled on 17^th^ to 22^nd^ day, the efficiency of the equipment remained at par with 9^th^ to 11^th^ day (65.8%). When bioacoustics was reinstalled for the second time at the experimental plot from 28^th^ to 30^th^, the efficiency of bioacoustics reduced to 41%. From 31^st^ to harvest of the crop, Parakeets did not visit the plot owing to reduced palatability (hardening) of achenes. Overall efficiency of the equipment was 71.6% for the entire experimental period.

### Experiment 13 Mar to 28 Mar 2014, Baswapur

During this experiment, the plot was damaged by Parakeets to an extent of 1.4% (negligible damage) before installing the equipment on13 Mar 2014. Distress feeding by Parakeets started on 10^th^ day of installing the bioacoustic equipment (Figure-6). Distress feeding happened for six days till the crop was harvested (29 Mar 2014).

From the land use and land cover maps for this experimental locality, we found that only 89% (69.42 km^2^) of the area had crops and trees providing food to Parakeets. Of this cropping extent, an average 68.9% of crops/trees bore food. It is imperative from (Table-6) that in the beginning of experiment, availability of food crops for Parakeets was plenty. However, on the date of distress feeding, availability of all of them had considerably decreased owing to harvest of crops.

**Table 6.** Extent of cropped area and availability of food to Parakeets, Baswapur, 13 Mar to 28 Mar 2014

| Crops with feeding stage for Parakeets in 78 km<sup>2</sup> | Extent of cropped area that has potential to provide food (ha) | Extent of area providing food as on 13 Mar 2014: beginning of experiment (ha) | Extent of area providing food as on 23 Mar 2014: at the start of distress feeding (ha) | Extent of area providing food as on 28 Mar 2014: at the harvest time (ha) |
| --- | --- | --- | --- | --- |
| Sunflower | 548 | 460 (84%) | 159 (29%) | 148 (27%) |
| Maize | 1173 | 1056 (90%) | 481 (41%) | 457 (39%) |
| Rabi Sorghum | 625 | 500 (80%) | 150 (24%) | 144 (23%) |
| Paddy | 1555 | 1135 (73%) | 358 (23%) | 342 (22%) |
| Red gram | 153 | 96 (63%) | 77 (50%) | 77 (50%) |
| Chickpea | 76 | 42 (55%) | 12 (16%) | 12 (16%) |
| Vegetables | 389 | 241 (62%) | 257 (66%) | 257 (66%) |
| Wild flowering and fruiting trees | 2186 | 984 (45%) | 1027 (47%) | 1027 (47%) |
| Orchards | 257 | 175 (68%) | 175 (68%) | 175 (68%) |

Visiting instances of Parakeets from the beginning of the experiment to 9^th^ day was 163/day. The instances of visits increased to 239/day from 10^th^ day till harvest. This showed that the surge in instance of Parakeets visiting experimental plots was correlated to shortage of food in the feeding areas. In this experiment, bioacoustics provided 89.6%% protection for first nine days in the absence of distress feeding. During the distress feeding stage, the efficiency of the equipment steeply decreased to 42% till the crop was harvested. Bioacoustics performed at an overall efficiency of 67.1%. Crop damage was negligible (1.4%) compared to 42% in control plot (Table-5).

## Experiment 20 Feb to 07 Mar 2013, IIMR

During this experiment, the Sorghum plot was damaged by Parakeets to an extent of 4.5% before installation of bioacoustics on 20 Feb 2013. Distress feeding by Parakeets started on 7^th^ day of installing the bioacoustic equipment (Figure-6). Distress feeding happened for nine days till the crop was harvested (08 Mar 2013).

From the land use and land cover maps for this experimental locality, we found that only 23.5% (18.33 km^2^) of the area had crops and trees providing food to Parakeets. This area was characterised by high percentage of urban structures. Of the cropping extent, an average 36.1% of crops/trees bore food at the beginning of the experiment. The area is characterised by typical small agricultural holdings fragmented by buildings. The date of sowing of all the crops differed to a great extent. Due to this, Parakeets constantly shifted feeding preferences depending on the maturity of food crops. Of all the available crops during the experiment, Sunflower was not available, but next highly preferred food, Sorghum was depredated. It is imperative from (Table-7) that in the beginning of experiment, availability of food crops for Parakeets was under short supply. However, on the date of distress feeding, availability of many of them had considerably decreased owing to harvest of surrounding crops in succession.

**Table 7.** Extent of cropped area and availability of food to Parakeets, IIMR, 13 Mar to 28 Mar 2013

| Crops with feeding stage for Parakeets in 78 km<sup>2</sup> | Extent of cropped area that has potential to provide food (ha) | Extent of area providing food as on 20 Feb 2013: beginning of experiment (ha) | Extent of area providing food as on 27 Feb 2013: at the start of distress feeding (ha) | Extent of area providing food as on 08 Mar 2013: at the harvest time (ha) |
| --- | --- | --- | --- | --- |
| Sunflower | 73 | 11 (15%) | Nil | Nil |
| Maize | 330 | 73 (22%) | 20 (6%) | 3 (1%) |
| Sorghum | 147 | 121 (82%) | 98 (67%) | 31 (21%) |
| Paddy | 422 | 274 (65%) | 219 (52%) | 186 (44%) |
| Red gram | 18 | 7 (40%) | 12 (65%) | 11 (61%) |
| Vegetables | 275 | 55 (20%) | 116 (42%) | 151 (55%) |
| Wild flowering and fruiting trees | 513 | 180 (35%) | 246 (48%) | 251 (49%) |
| Orchards | 55 | 6 (10%) | 7 (13%) | 8 (14%) |

Visiting instances of Parakeets from the beginning of the experiment to 6^th^ day was 113/day. The instances of visits increased to 1321/day from 7^th^ day till harvest. This showed that the surge in instance of Parakeets visiting experimental plots was correlated to shortage of food in the feeding areas. In this experiment, bioacoustics provided 89.6%% protection for first nine days in the absence of distress feeding. During the distress feeding stage, the efficiency of the equipment steeply decreased to 42% till the crop was harvested. Bioacoustics performed at an overall efficiency of 67.1%. Crop damage was negligible (1.4%) compared to 42% in control plot (Table-5).

## Conclusions

- Distress feeding almost always resulted in sudden surge of visiting instances by Parakeets independent of crop types. There was no surge in instances of Baya Weavers visiting the field during distress feeding. This was due to very limited and localised feeding behaviour of the breeding birds, apart from absence of choice of food based on palatability by Bayas.
- There was a strong correlation between distress feeding and availability of food in the established feeding range of Parakeets.
- Bioacoustics protected the crops during distress feeding, albeit at a reduced percentage. Removal of bioacoustics during distress feeding resulted into increase in visiting instances by Parakeets. The surge in percentage of Parakeet visits to experimental fields during distress feeding ranged from 47% to 3025%.
- Distress feeding in isolated situations (Figure-6) leads to huge surge in visiting instances of Parakeets resulting into total loss of crops. Nearly all reported cases of total devastation of crops by depredatory birds are always due to distress feeding.
- Distress feeding is a function of food preference (palatability), extent of cropping area, roost location, physiological condition of the bird, and breeding season.
- Distress feeding can be avoided by synchronized sowing and increasing the cropping extent

## Acknowledgement

We thank National Agricultural Science Foundation (NASF), ICAR, New Delhi, for funding this project. We also extend our thanks to Professor Jayashankar Telangana State Agriculture University, Hyderabad for providing required facilities to conduct experiments.

## References

AINPAO. (2000). Research Accomplishments Agricultural Ornithology Technical Bulletin-II, All India Network Project on Agricultural Ornithology, ANGRAU, Rajendranagar, Hyderabad. 37.

AINPAO. (2012). Annual report of Agricultural Ornithology, All India Network Project on Agricultural Ornithology, ANGRAU, Rajendranagar, Hyderabad. 37.

Fitzgerald, S. (2013). Managing bird damage in crops. Fruit and vegetable gardener’s association, Ontario, 6.

Greene, T.C. (1988). Foraging ecology of the Red-crowned Parakeet and Yellow-crowned Parakeet on Little Barrier Island, Hauraki Gulf, New Zealand. New Zealand Journal of Ecology. 22(2): 161–171.

Institute for Land, Water and Society. (2013). Managing agricultural landscapes to maximise production and conservation outcomes: The case of the Regent Parrot. Final report, Australian Research Council Linkage Grant. 21.

Imadullah, M. (2014). Time series analysis. Basic statistics and data analysis. itfeature.com

Likert, R. (1932). A technique for the measurement of attitudes. Archives of Psychology. 140: 1–55.

Pestgo Airport Wailer. (2017). Pestgo Airport wailer MkIII, Hughes and Hughes, UK, http://www.hugheschem.com/ProductDetail.aspx?productId=11&cvalu=Airport&ctype1=NA&proid=NA

Philip, C. Whitford. (2009). Succesful use of alarm and alert calls to reduce emerging crop damage by resident Canada Geese near Horizon Marsh, Wisconsin. Biology Department, Capital University, Columbus, Ohio. Bird strike North America conference, Paper-1. 74– 79

Surender, G. and Swamy, K. (2014). Pers. Comm. On feeding of partially maimed Rose-ringed Parakeets on bioacoustics protected sunflower crop, Sira, Karnataka. Jan 2014.

Upton, G. and Cook, J. (1996). Understanding Statistics. Oxford University press, 55.

